# Using Diffusion Transformers to Generate Synthetic Diffusion Scalar Maps for Data Augmentation

**DOI:** 10.1101/2025.08.27.672680

**Authors:** Tamoghna Chattopadhyay, Chirag Jagad, Pavithra Senthilkumar, Sophia I. Thomopoulos, Julio E. Villalón-Reina, Paul M. Thompson

## Abstract

Generation of high-quality synthetic brain MRI data could be beneficial for advancing neuroimaging research, particularly when access to large-scale, labeled datasets is limited. In this work, we leverage a pretrained Diffusion Transformer (DiT) architecture to synthesize 3D mean diffusivity (MD) scalar maps from the Cam-CAN dataset. To adapt the DiT model—originally trained on 2D natural images—for 3D neuroimaging data, we implemented a preprocessing strategy that tiles 2D slices from 3D volumes into composite 2D images, enabling effective finetuning. The quality of the generated synthetic images was evaluated using Multi-Scale Structural Similarity (MS-SSIM) and Maximum Mean Discrepancy (MMD) metrics, demonstrating high fidelity and anatomical coherence. To assess the utility of synthetic data in downstream tasks, we conducted transfer learning experiments for dementia classification on the ADNI dataset. A sex classification model, trained on both real and synthetic Cam-CAN data, was repurposed for this task, showing that synthetic samples can enhance model performance. These results highlight the potential of diffusion-based generative models for augmenting neuroimaging datasets and supporting clinical applications.

## I. Introduction

The acquisition of high-resolution diffusion MRI (dMRI) data, particularly Diffusion Tensor Imaging (DTI [1]) scalar maps including mean diffusivity (MD), remains resource-intensive due to relatively long scan times, sensitivity to motion artifacts, and complex preprocessing workflows. Synthetic data generation offers a pathway to address data scarcity and privacy concerns, yet existing methods struggle to provide anatomical fidelity, computational efficiency, and compatibility with downstream clinical tasks. Recent advances in diffusion models and transformer architectures present new opportunities for cross-modal image synthesis, yet innovative solutions are needed to adapt these frameworks to 3D neuroimaging-particularly for translating structural T1-weighted (T1w) MRI to diffusion MRI metrics..

Traditional approaches using Generative Adversarial Networks (GANs [2,3]) have enabled T1w-to-T2w translation and slice-wise dMRI synthesis. However, methods like CycleGAN often fail to preserve microstructural features critical for diffusion scalars, while 3D GANs face memory constraints and training instability. Gu et al. [4] used a 2D CycleGAN to generate FA and MD map slices from T1w slices. Pan et al. [5] used a transformer-based encoder along with conditional GANs to translate 2D T1w to T2w slices. Chattopadhyay et al. [6,7] used denoising diffusion probabilistic models (DDPMs) and latent diffusion models (LDMs) to generate 2D and 3D DTI-MD maps from T1w MRIs, though they were not one-to-one mapped. Zhang et al. [8] used a diffusion bridge model to generate 3D DTI maps from T1w MRI scans. In contrast, Diffusion Transformers (DiTs [9]) combine the probabilistic framework of diffusion models with transformer scalability, achieving state-of-the-art results in natural image synthesis. They represent a significant advancement in generative AI, merging the strengths of transformers and diffusion models, enabling faster training and improved scalability. Li et al. [10] introduced MedDiT, a knowledge-controlled diffusion transformer framework that generates plausible medical images aligned with simulated patient symptoms. Even so, there are few studies evaluating DiTs for cross-modal MRI synthesis-specifically for translating structural T1-weighted scans to diffusion-derived scalar maps.

In this work, we address these limitations by adapting a pretrained 2D DiT model to synthesize 3D MD maps from the Cam-CAN [11] dataset. We introduce a novel preprocessing strategy that tiles 2D slices from 3D volumes into composite images, enabling efficient finetuning without sacrificing anatomical consistency. We evaluate synthetic image quality using Multi-Scale Structural Similarity (MS-SSIM) [12] and Maximum Mean Discrepancy (MMD) [13], ensuring pixel-level accuracy and distributional alignment with real data. To demonstrate clinical relevance, we repurpose a sex classification model trained on hybrid (real + synthetic) data for dementia classification on the Alzheimer’s Disease Neuroimaging Initiative (ADNI [14]) dataset, showcasing the utility of synthetic data in transfer learning scenarios.

Our approach introduces two innovations:

1. M**emory-efficient preprocessing:** 3D volumes are tiled into composite 2D slices, enabling finetuning of DiT-originally trained on natural images-without architectural modifications.
2. **Downstream-task optimization:** Synthetic data is validated not only via traditional metrics (MS-SSIM, MMD) but also through transfer learning performance in dementia classification, addressing limitations of prior evaluations that focused solely on image quality.

## II. Imaging Data and Preprocessing

The primary dataset for our experiments was the Cambridge Centre for Aging and Neuroscience (Cam-CAN) dataset [11], chosen here for its broad age distribution (range: 18-88 years). For the downstream task, we also used the Alzheimer’s Disease Neuroimaging Initiative (ADNI) dataset – a multisite study launched in 2004 to improve clinical trials to prevent and treat AD [14]. The data distribution is shown in **Table 1**; we divided the ADNI dataset into train, validation and test sets in the ratio 80:10:10. We ensured that there was no overlap between the testing and training data subsets for downstream task, and that the test dataset had only one scan per subject. 3D T1-weighted (T1w) brain MRI volumes were pre-processed using the following steps [15]: nonparametric intensity normalization (N4 bias field correction), ‘skull stripping’ for brain extraction, nonlinear registration to an in-house template [15] with 6 degrees of freedom and isometric voxel resampling to 2 mm. The pre-processed images were of size 91×109×91. The T1w images were scaled using min-max scaling to take values between 0 and 1. Images were resized to 64×64×64 to fit the DiT architecture. All T1w images were aligned to the ENIGMA template provided by the ENIGMA consortium [16].

**Table 1.** Data distribution for experiments.

| Dataset | Total N | Sex | | Mean Age ± St. Dev. | Diagnosis | |
| --- | --- | --- | --- | --- | --- | --- |
|  |  | Male | Female |  | CN | Dementia |
| Cam-CAN | 652 | 330 | 322 | $54.30 \pm 18.60$ | 652 | - |
| ADNI | 332 | 169 | 163 | $74.61 \pm 8.14$ | 199 | 133 |

## III. Deep Learning Architectures

Our framework builds on the Latent Diffusion Transformer (DiT) architecture DiT-XL/2 [9], which replaces traditional U-Net [17] backbones with transformer blocks [18] for noise prediction. The overall pipeline involves compressing high-dimensional input images or 3D volumes into a compact latent representation using a pre-trained Variational Autoencoder (VAE [19]). For 2D images of size 512×512×3, the VAE reduces the spatial dimensions by a factor of 8, resulting in a latent space of 64×64×4. In the case of 3D volumes, we first uniformly resize each sample to 64×64×64, then decompose them along one axis into 64 slices to form a 2D grid of images. These slices are treated as independent samples for training and inference. The latent representations are partitioned into non-overlapping square patches of size *p*×*p*, which are then flattened and projected into a fixed-dimensional embedding space. This sequence of patch embeddings is processed by a series of transformer blocks, each consisting of multi-head self-attention layers and feed-forward modules. Positional information is preserved by adding sinusoidal positional encodings to each patch embedding.

A key innovation in DiT is the use of in-context conditioning, where the diffusion timestep *t* and class label *c* modulate both the adaptive layer normalization and the pre-activation scaling layers within each transformer block. This design enhances the model’s flexibility, so it can adapt dynamically during the denoising process. We adopt the largest available DiT variant, DiT-XL/2, which features 28 transformer layers, each with 16 attention heads and a hidden dimension of 1,152. This configuration offers strong modeling capacity while maintaining a fixed number of parameters regardless of the image resolution. Empirical results from prior work indicate that this model size, particularly with a patch size of *p* = 2, yields optimal image generation quality. We employed the Adam optimizer with a learning rate of 5e-4 and used mean squared error (MSE) as the loss function. Given the input image resolution of 512×512, the corresponding latent representation had dimensions of 64×64. During fine-tuning, only the bias terms, layer normalization parameters, and class embeddings were updated, while all other weights were kept frozen. Sex was used as the class for training, with 0/1 for Male/Female. After finetuning the DiT-XL/2 model on decomposed 2D slices, we generate synthetic 2D image grids. These are subsequently reassembled by stacking the generated slices to reconstruct the full 3D volumes, enabling the synthesis of high-fidelity 3D data from 2D representations. To evaluate the quality of the generated synthetic images, we adopted standard metrics commonly used in generative model assessment. Specifically, we measured both the realism and diversity of the outputs across 500 image pairs using MMD and MS-SSIM. A lower MMD score suggests a closer alignment between the distributions of synthetic and real images, while a higher MS-SSIM score indicates stronger structural resemblance.

To evaluate realism, comparisons were made between real and synthetic images from the same class using both MMD and MS-SSIM. To assess diversity, MS-SSIM was computed between pairs of synthetic images generated under identical class conditions, with comparisons to real image pairs providing a reference baseline.

The 3D CNN model architecture (**Fig. 2**) comprised four convolutional layers with 3×3 kernels, followed by a 1×1 convolution and a final fully connected layer. ReLU activation and Instance Normalization were applied throughout to enhance training stability. To address overfitting, dropout layers with a 0.5 rate were added after the second, third, and fourth convolutional layers, along with a 3D average pooling layer using a 2×2 kernel. The model was trained with the Adam optimizer [20], initialized with a learning rate of 1×10^−4^ and an exponential decay factor of 0.96. Training was performed over 100 epochs with a batch size of 8, optimizing for mean squared error. Early stopping and dropout were used as regularization strategies. Hyperparameters were tuned to identify the optimal configuration. We performed model evaluation using model weights from sex classification (a standard benchmark with known ground truth) as pretraining for dementia classification, using balanced accuracy and F1 score as performance metrics. We compared model performance after pretraining on different combinations of original and synthetic DTI-MD maps.

**Figure 1.**
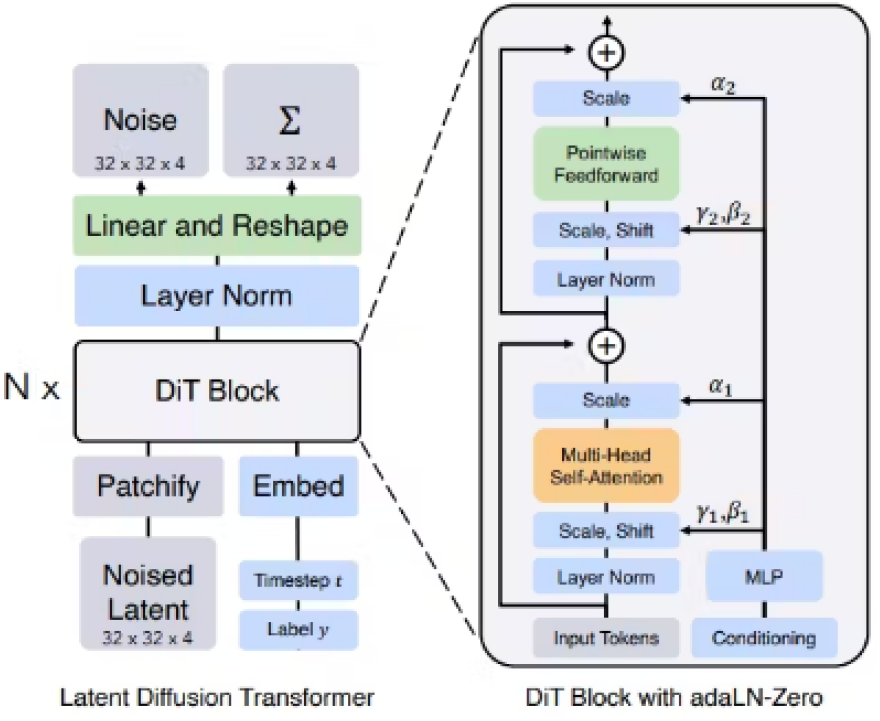
DiT architecture, reproduced from [9].

**Figure 2.**
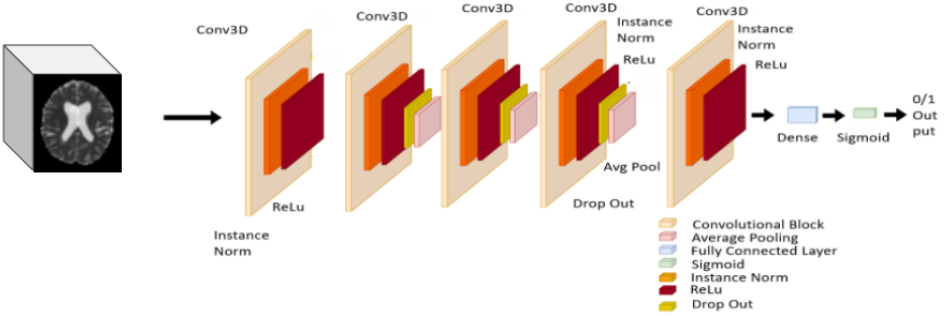
3D CNN Architecture that we trained for downstream tasks.

**Figure 3.**
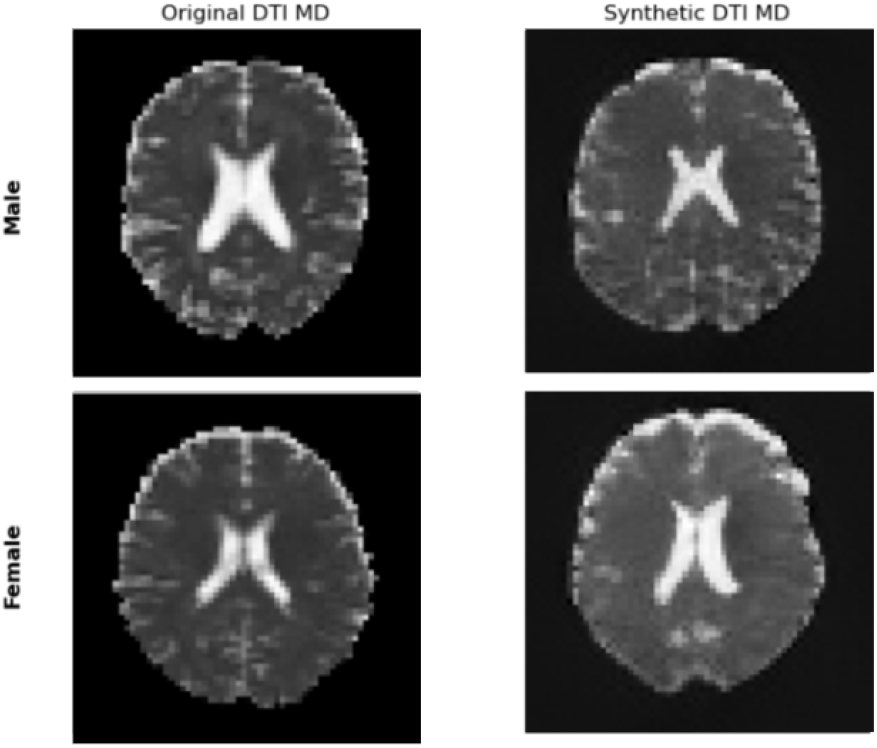
Results from the DiT implementation. Each column shows the middle slice from the original DTI-MD scan and the synthetic DTI-MD scan respectively. The first row is for a male subject and second row is for a female subject.

## IV. Results

Evaluations of the synthetic DTI-MD maps generated by the DiT model, compared to real scans, are summarized in **Table 2**. For both male and female participants, the DiT-based model demonstrated strong performance across realism and diversity metrics. In terms of realism, the Maximum Mean Discrepancy (MMD) scores indicate a low statistical distance between the synthetic and real distributions. Additionally, the MS-SSIM (Real vs. Synthetic) values used to assess structural similarity suggest good anatomical fidelity, with a slight edge observed in the female cohort. In terms of diversity, MS-SSIM (Real vs. Real) scores provide a benchmark for expected variability in the real data. The diversity among the synthetic samples themselves, is measured via MS-SSIM (Synthetic vs. Synthetic). The high scores reflect substantial intra-group variability, suggesting that the DiT model generates a broad and heterogeneous set of synthetic images. Overall, these results indicate that the DiT-based generative model effectively balances anatomical realism and sample diversity. The consistently high diversity scores support its capacity to capture a wide range of anatomical patterns, while the favorable realism metrics show that it can generate structurally coherent, biologically plausible data. This balance is important for downstream tasks such as transfer learning or data augmentation in clinical neuroimaging applications.

**Table 2.** Qualitative Metrics for the Synthetic Dataset.

|  | Male | Female |
| --- | --- | --- |
| **Epochs** | 2,000 | 2,000 |
| **Image Size** | 64x64x64 | 64x64x64 |
| **Realism: MMD (Real vs Synthetic)↓** | $0.0275 \pm 0.0073$ | $0.0238 \pm 0.0065$ |
| **Realism: MS-SSIM (Real vs Synthetic)↑** | $0.5299 \pm 0.1704$ | $0.6251 \pm 0.1295$ |
| **Diversity: MS-SSIM (Synthetic vs Synthetic)↓** | $0.6522 \pm 0.2375$ | $0.6974 \pm 0.1879$ |
| **Diversity: MS-SSIM (Real vs Real)↓** | $0.8762 \pm 0.0239$ | $0.8864 \pm 0.0151$ |

To assess the clinical relevance of the generated synthetic data, we repurposed a convolutional neural network (CNN) model, initially trained for sex classification on the Cam-CAN dataset, for downstream dementia classification using the ADNI dataset. This experiment evaluates the utility of synthetic data in transfer learning settings, where pretrained model weights are fine-tuned for a related clinical task.

We report results using balanced accuracy and F1 score, averaged over five independent runs (**Table 3**). Increasing the volume of synthetic data during the pretraining task enhanced generalization for the downstream clinical task. In contrast, pretraining on only synthetic data (0% real, 100% synthetic) resulted in poorer performance, highlighting the importance of including real data during pretraining. These results show that synthetic data, when combined with real data, can effectively boost downstream performance and support clinical applications through improved model robustness and transferability.

**Table 3.** Classification results for the downstream task. B.A. indicates balanced accuracy and F1 indicates F1 Score. The percentages for synthetic data are expressed as a fraction of the total amount of real data.

| Experimental Setup | B.A. | F1 |
| --- | --- | --- |
| Pretrained on Real 100%, Synthetic 0% | 0.780 | 0.713 |
| Pretrained on Real 100%, Synthetic 100% | 0.781 | 0.716 |
| Pretrained on Real 100%, Synthetic 200% | **0.803** | **0.745** |
| Pretrained on Real 0%, Synthetic 100% | 0.756 | 0.681 |

## V. Conclusions and Future Work

In this study, we evaluated the use of a Diffusion Transformer architecture for synthesizing mean diffusivity maps from structural T1-weighted MRI, leveraging the Cam-CAN dataset and evaluating clinical relevance through transfer learning on the ADNI dataset. By adapting DiT for 3D neuroimaging through a novel 2D tiling strategy, we achieved strong performance on both realism and diversity metrics. In quantitative evaluations, the synthetic images closely resembled real MD maps, with low MMD and high MS-SSIM scores across both male and female subgroups. Diversity metrics further confirmed that the synthetic data captured a wide range of anatomical variability, critical for building robust machine learning models. To assess clinical utility, we repurposed a CNN model trained on real and synthetic data for downstream dementia classification. Inclusion of synthetic samples improved performance in a dose-dependent manner, with models trained on real data augmented by synthetic MD maps outperforming those trained on real data alone. These findings affirm the potential of DiT-generated synthetic diffusion MRI for enhancing model generalization and addressing data scarcity in neuroimaging. Future work will examine architectural improvements for fully 3D diffusion-based models. Expanding to multi-channel diffusion synthesis—such as fractional anisotropy (FA), radial diffusivity (RD), and axial diffusivity (AD)—is also a key direction, enabling richer feature representation for clinical prediction tasks.

## Acknowledgments

This work was supported by NIH/NIA grant U01 AG068057 (‘AI4AD’).

